# Pennycress (*Thlaspi arvense* L.) Seed Persistence in the Field After Two Years

**DOI:** 10.1101/2025.04.18.649506

**Authors:** Tad L. Wesley, Mary E. Phippen, Winthrop B. Phippen

**Affiliations:** School of Agriculture, Western Illinois University, Macomb, IL, USA

## Abstract

Field pennycress (*Thlaspi arvense* L.), is a new oilseed winter annual crop being investigated as a source of biofuel in the United States. The purpose of this study is to assess long term survivability and dormancy of both wild-type pennycress and gene-edited golden pennycress seed. Seeds from one wild type and three gene-edited golden types that carried an edit to *TT8* gene were buried at 2 cm and 15 cm depths under well-drained and poorly-drained field conditions. The seeds were exhumed at 2, 4, 6, 12, 18, and 24 months after burial and germinated to determine the viability of the seed over time. All golden seeded gene-edited lines decreased to 0% germination by 6 months while the wild black seeded variety ‘ARV1’ retained almost 60% germination after 2 years. A significant difference was seen in the ‘ARV1’ survival in the well-drained field but not in the poorly-drained field. However, there was no significant difference in seed viability for burial depth in the well-drained field but there was significant difference for burial depth in the poorly-drained field. These results indicate that gold seeded varieties carrying the edit to *TT8* through gene-editing have dramatically decreased the survivability of the seed in the seedbank. Reduced survivability will greatly assist in the adoption of golden pennycress as a new viable off-season crop in the Midwest without concerns of adding to the seedbank or serving as weed pressure in primary crops.

## Introduction

Pennycress, (*Thlaspi arvense* L.), a member of the Brassicaceae family closely related to rapeseed, canola, carinata, and camelina, is an intermediate oilseed energy crop well suited for cultivation in the U.S. Midwest (Phippen et al., 2022; Basnet and Ellison, 2024). With extreme winter hardiness, high oilseed yields, and a short life cycle, pennycress fits between primary crops within the 80 million-acre Midwest Corn Belt (Phippen and Phippen, 2012; Fan et al., 2013; Johnson et al., 2015; Ott et al., 2019). Cultivating pennycress during fallow periods avoids direct and indirect land-use changes typically associated with agricultural intensification (Fargione et al., 2008; Searchinger et al., 2008). In addition, pennycress provides the ecosystem benefits of a cover crop while generating net profit for producers (Plastina et al., 2018; Shi et al., 2019).

Pennycress seeds contain ∼35% oil with a fatty acid profile suitable for biofuels production, meeting the U.S. Renewable Fuels Standard (Moser et al., 2009; Moser, 2012). Using CRISPR genome editing, pennycress oil composition has been further improved by eliminating erucic acid (C22:1) content and increasing oleic acid (C18:1) content through knocking out Fatty Acid Elongase 1 (*FAE1*); thereby making an oil equivalent in composition to canola and improving efficiency of its conversion to Sustainable Aviation Fuel (SAF) and other biofuels (McGinn et al., 2019; Chopra et al., 2020; Jarvis et al., 2021). In addition, the discovery of knocking out the Transparent Testa 8 (*TT8*) gene has been a critical step in helping to domesticate pennycress (Ulmasov et al., 2020). *TT8* is a transcription factor acting as a master regulator of seed coat formation (Chopra et al., 2018). Wild pennycress seeds have thick seed coats conferring dormancy and prolonged protection against the elements. Knockout of *TT8* eliminates these weediness traits, reducing seed coat thickness and condensed tannins, resulting in golden colored (“golden pennycress”) seeds that germinate much better than wild type, dramatically improving crop establishment. Reducing the seed coat thickness has also been shown to reduce seed persistence in the soil (Gardarin et al., 2010). *TT8* and *FAE1* knockout mutations do not impact plant health and form the basis of all commercial pennycress varieties (CoverCress®).

One obstacle to pennycress implementation throughout the entire US Midwest region is that wild pennycress is known to persist in cultivated soil for up to 6 years (Chepil, 1946) and undisturbed soils for 17 to 30 years (Burnside, 1996). Baskin and Baskin (1989) have shown seeds of winter annual plants of pennycress are non-dormant in autumn and enter dormancy in winter months under cooler temperatures. Seed longevity is not the only factor contributing to the seriousness of a pennycress weed problem. An important factor in the persistence of pennycress may be the tremendously large numbers of seeds produced annually by each plant. Mitich (1996) reported that wild pennycress plants produce 1,600 to 50,000 seeds per plant during a growing season. The survival to maturity of only a few seeds of pennycress each year would be sufficient to re-infest the area with several thousand fresh seeds.

The purpose of this study is to address the concerns of possibly spreading or increasing weed pressures on primary crops due to commercial pennycress production. The main objective of this study was to bury a wild type black seeded pennycress line and 3 gene-edited golden lines to study seed survivability over a 2-year period in well-drained and poorly-drained field conditions at burial depths of 2 cm and 15 cm.

## Materials and method

### Seed source

Mature seed from one wild population, one gene-edited golden variety, and two commercial golden gene-edited lines was collected in late spring of 2022 from mature research plots at WIU in Macomb, IL (40°27′38″ N, 90°40′27″ W). Varieties included: a wild type black seeded parental breeding line ‘ARV1’, a golden seeded gene edited variety ‘tt8-t/ARV1’ in the ‘ARV1’ background, and 2 commercial golden seeded varieties identified as ‘CC1’ and ‘CC2’ with similar *TT8* edits. All seed was tested at time 0 and produced germination rates above 95%.

### Burial

One hundred seed of each line were mixed with a silt loam soil collected from the burial location and placed in 5 cm x 7 cm polyester mesh bags with 125-µm mesh openings (one variety per bag), and buried at 2 and 15 cm on June 15, 2022 at the WIU Research Farm in Macomb, IL. Bags were laid flat at each depth in 2 areas of a no-till corn and soy production field. The first area represents a well-drained field, Greenbush silt loam, while the second area represents a poorly-drained location of the field Sable silty clay loam. Bags were tied to a flag with nylon filament at each location. Bags (four replications for each variety and burial depth) were exhumed 2, 4, 6, 12, 18, and 24 months after burial. Overall, there were 6 exhume times with 2 depths and 2 drainages for each variety.

### Seed germination

Immediately after the bags were exhumed from the field conditions, bags were washed in running tap water to remove silt. Intact seeds were removed from the bags, placed on moist blue blotter germination paper, and incubated for 21 days at 23°C. Seeds that germinated were counted and removed at 7, 14, and 21 days. Germinated seeds were distinguished by having both a shoot and a radicle emerge from the seed. After 21 days, intact un-germinated seeds were placed in a 3,000 ppm 2,3,5-triphenyltetrazolium chloride (TZ) solution (pH 7.0) for 7 days (Figure 1). Seeds were dissected under a microscope and seeds with dark pink to red embryos were recorded as viable (Figure 2). All other seeds were recorded as non-viable.

**Figure 1.**
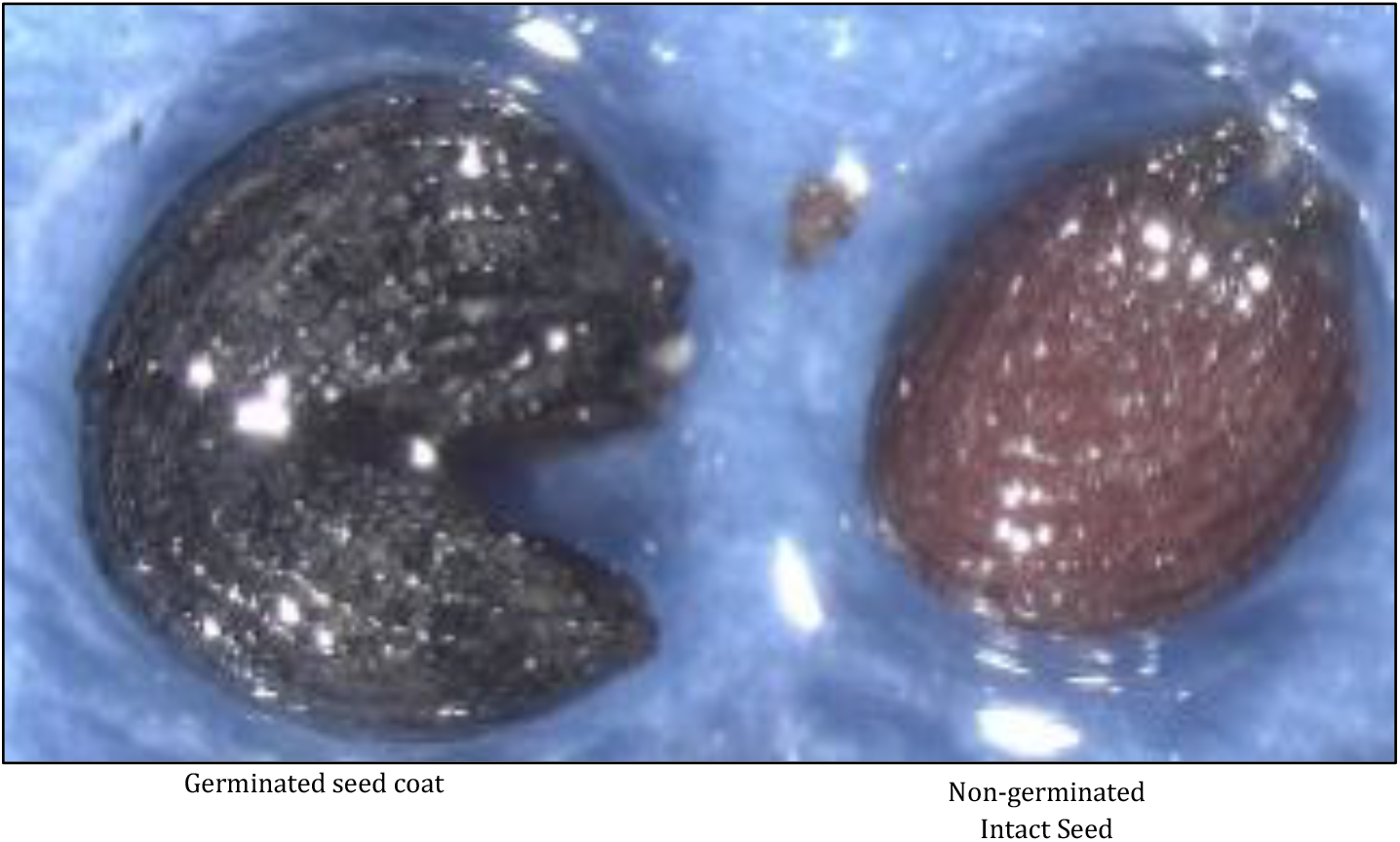
non-germinated seed next to germinated seed coat.

**Figure 2.**
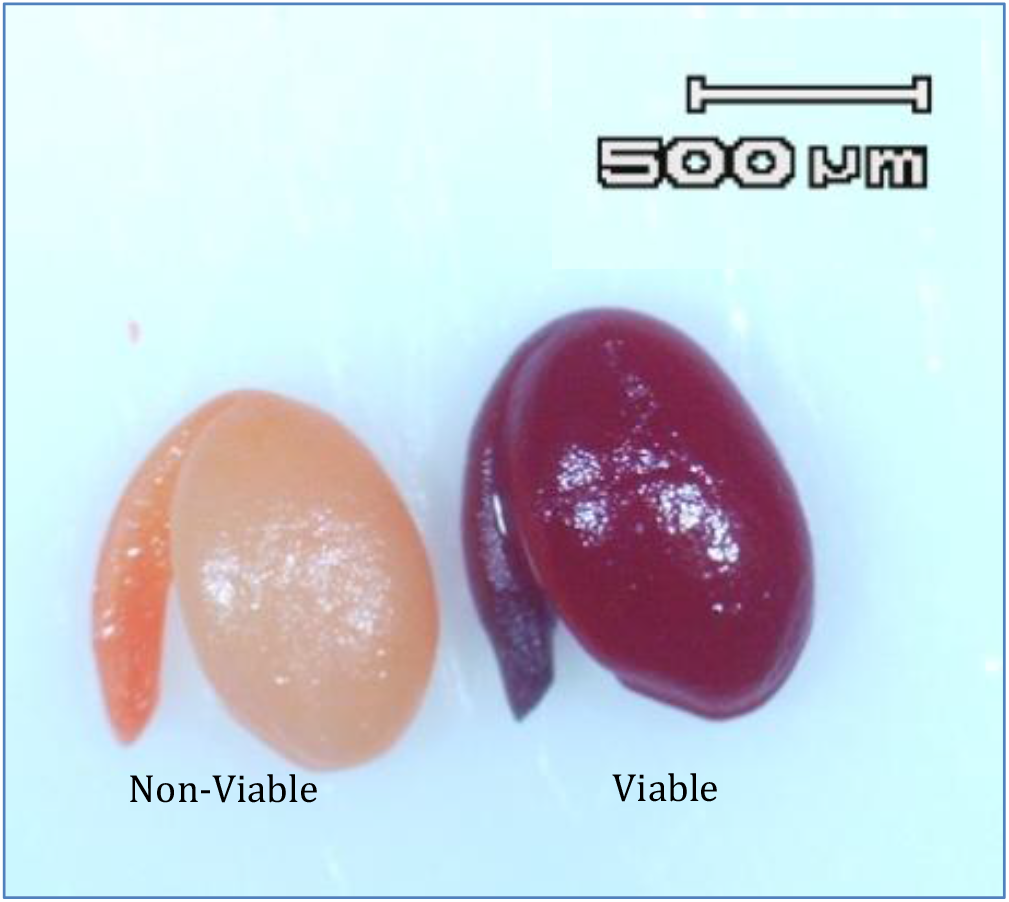
Comparison of viable and non-viable seed after TZ test.

### Data analysis

RStudio was used to statistically analyze all data and produce the graphs. For germination and viability data collected at 24 months, summary statistics (mean, standard deviation, minimum, and maximum values) were calculated. An ANOVA (Analysis of Variance) was performed to assess the effects of soil drainage (well drained and poorly drained) and burial depth (2 cm and 15 cm) on germination rates and seed viability. Pairwise comparisons were conducted using Tukey’s HSD post-hoc test at a 95% confidence level to determine statistically significant differences among treatments. The time-dependent relationships for germination rates and viability for seed buried in both poorly and well-drained soil types at both 2 cm and 15 cm were investigated. All weather data throughout the study were collected from an on-site weather station and 10-year averages were collected from the Macomb, Illinois Airport data archived by the Illinois Climate Network (2023).

## Results and discussion

The golden pennycress had low germination rates and no viable seed by 2 months after burial and no germination or viable seed at 12 months of burial as seen in Figures 3 and 4. This dramatic reduction in viability is likely associated with the thin golden seed coat. Whereas, the average germination rates of wild thick seed coat black pennycress ‘ARV1’ were 80% after 2 months and declined to only 50% after 2 years of burial across all treatments. The thick double cell layer of the black seed coat helps the seed remain impermeable to water and maintain seed longevity. After 2 years, all of the non-germinated wild ‘ARV1’ pennycress was still viable after 21 days in the incubator for both soil type locations (Table 1). The 2 cm burial depth in the well-drained location was significantly different from all other burial depths and locations for black seed germination (Figure 5). Solarization, rapid heating and cooling at the soil surface is known to help diminish weed seeds in production fields (Patell, 2005). Pennycress seeds at the shallow 2 cm depth would have experienced dramatic fluctuations in soil moisture and temperature throughout the summer and fall months leading to a rapid decline in viability. It has been shown that black wild pennycress seed longevity is improved by storage under cooler temperatures of 4°C and -34°C for up to 2 years. However, black seeds stored at room temperature (22°C) had significantly decreased germination rates (Brandhorst et al., 2024). Black seeds buried at 15 cm in poorly drained soils would have had a cooler and more stable environment that could improve seed longevity.

**Table 1.**
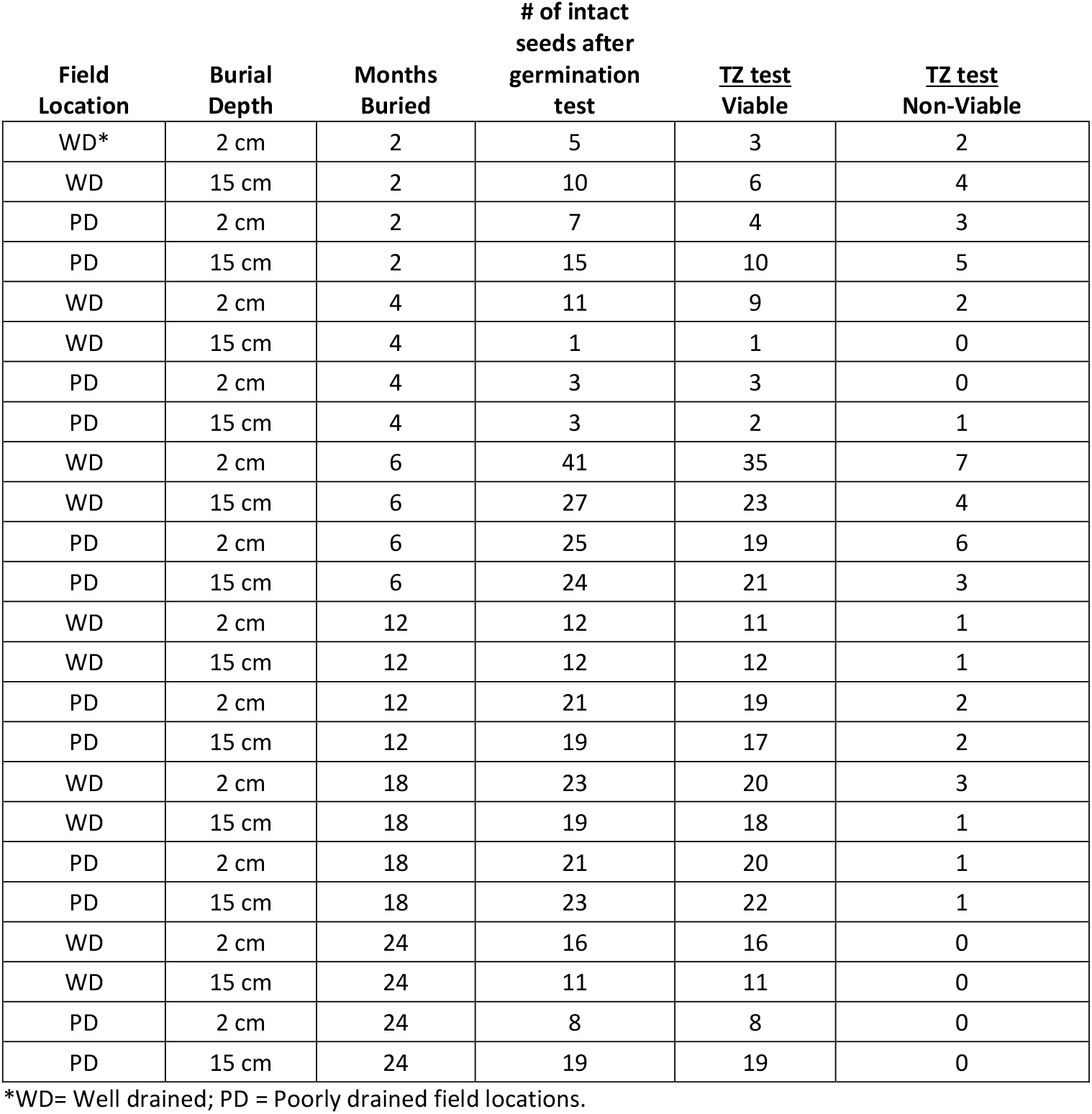
Tetrazolium seed viability results for all un-germinated black seeded ‘ARV1’ seeds after 24 months of burial.

| Field Location | Burial Depth | Months Buried | # of intact seeds after germination test | <u>TZ test</u><br>Viable | <u>TZ test</u><br>Non-Viable |
| --- | --- | --- | --- | --- | --- |
| WD* | 2 cm | 2 | 5 | 3 | 2 |
| WD | 15 cm | 2 | 10 | 6 | 4 |
| PD | 2 cm | 2 | 7 | 4 | 3 |
| PD | 15 cm | 2 | 15 | 10 | 5 |
| WD | 2 cm | 4 | 11 | 9 | 2 |
| WD | 15 cm | 4 | 1 | 1 | 0 |
| PD | 2 cm | 4 | 3 | 3 | 0 |
| PD | 15 cm | 4 | 3 | 2 | 1 |
| WD | 2 cm | 6 | 41 | 35 | 7 |
| WD | 15 cm | 6 | 27 | 23 | 4 |
| PD | 2 cm | 6 | 25 | 19 | 6 |
| PD | 15 cm | 6 | 24 | 21 | 3 |
| WD | 2 cm | 12 | 12 | 11 | 1 |
| WD | 15 cm | 12 | 12 | 12 | 1 |
| PD | 2 cm | 12 | 21 | 19 | 2 |
| PD | 15 cm | 12 | 19 | 17 | 2 |
| WD | 2 cm | 18 | 23 | 20 | 3 |
| WD | 15 cm | 18 | 19 | 18 | 1 |
| PD | 2 cm | 18 | 21 | 20 | 1 |
| PD | 15 cm | 18 | 23 | 22 | 1 |
| WD | 2 cm | 24 | 16 | 16 | 0 |
| WD | 15 cm | 24 | 11 | 11 | 0 |
| PD | 2 cm | 24 | 8 | 8 | 0 |
| PD | 15 cm | 24 | 19 | 19 | 0 |

\*WD= Well drained; PD = Poorly drained field locations.

**Figure 3.**
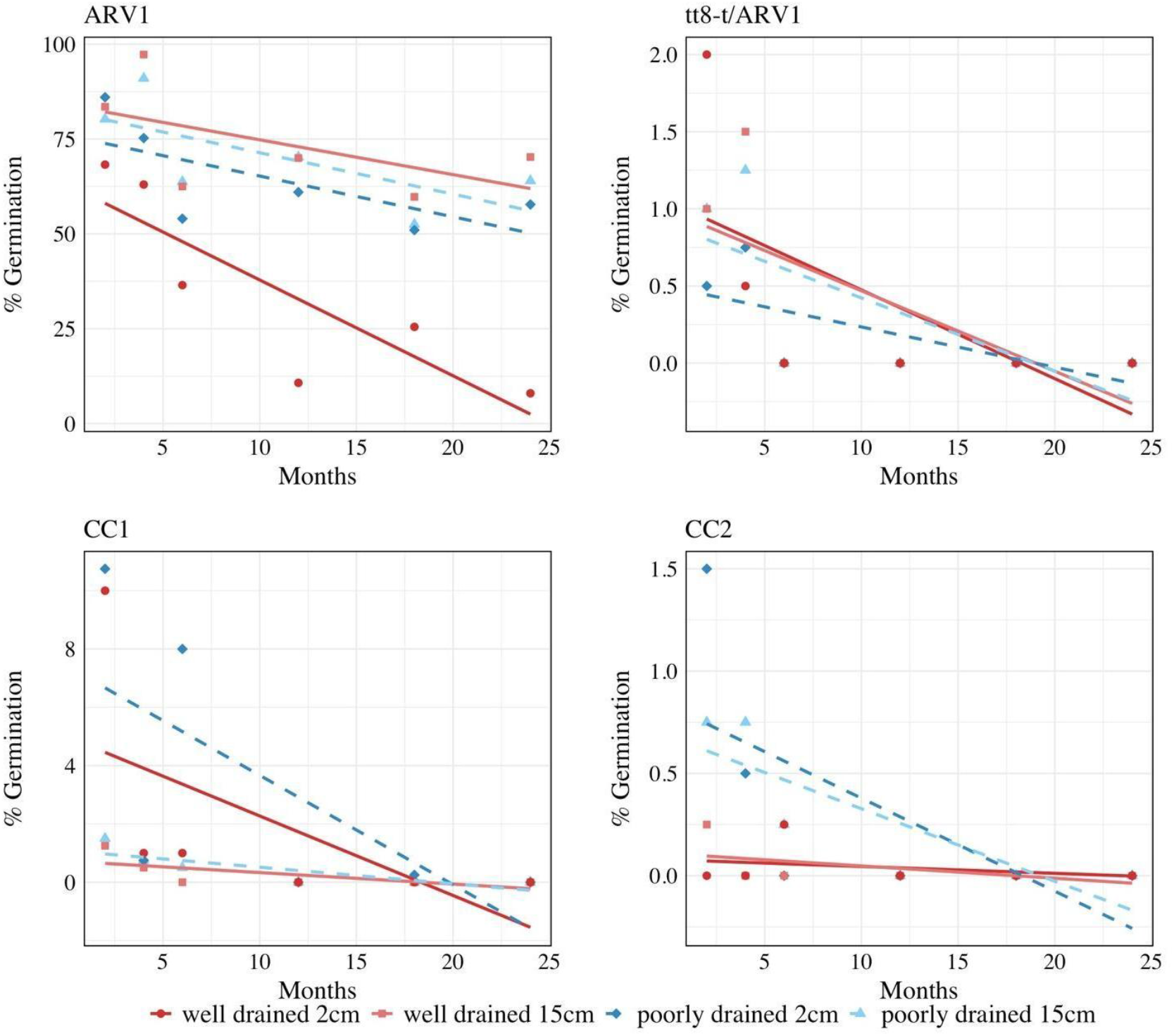
Twenty-one day percentage seed germination results after 24 months of seed burial for all seed varieties. Data represents averaged seed germination for all locations and burial depths (‘ARV1’ is black seeded, while ‘CC1’, ‘CC2’, and ‘tt8-t/ARV1’ are all gene-edited golden seeded).

**Figure 4.**
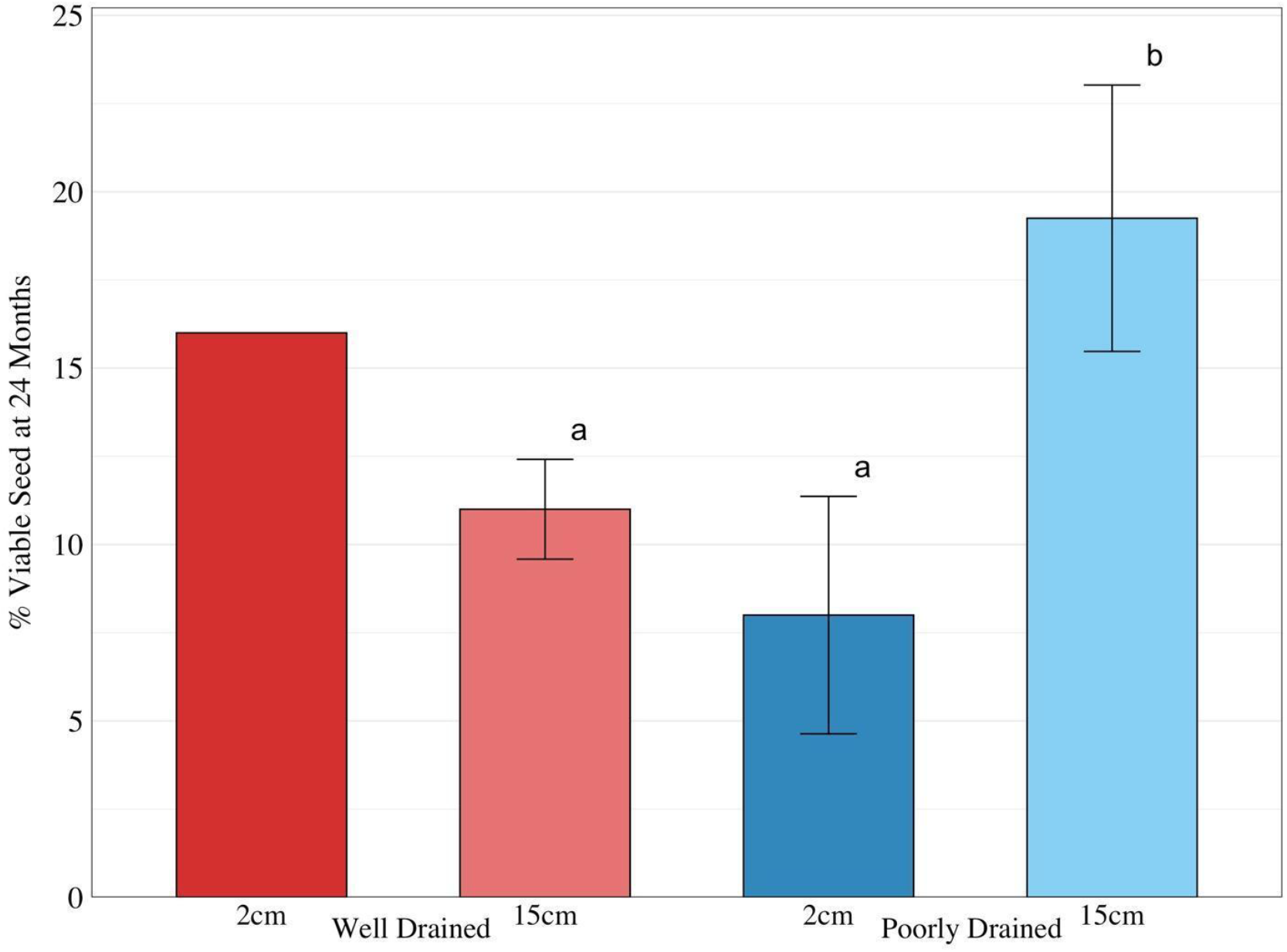
Tetrazolium seed viability results for un-germinated ‘ARV1’ seeds after 24 months of burial.

**Figure 5.**
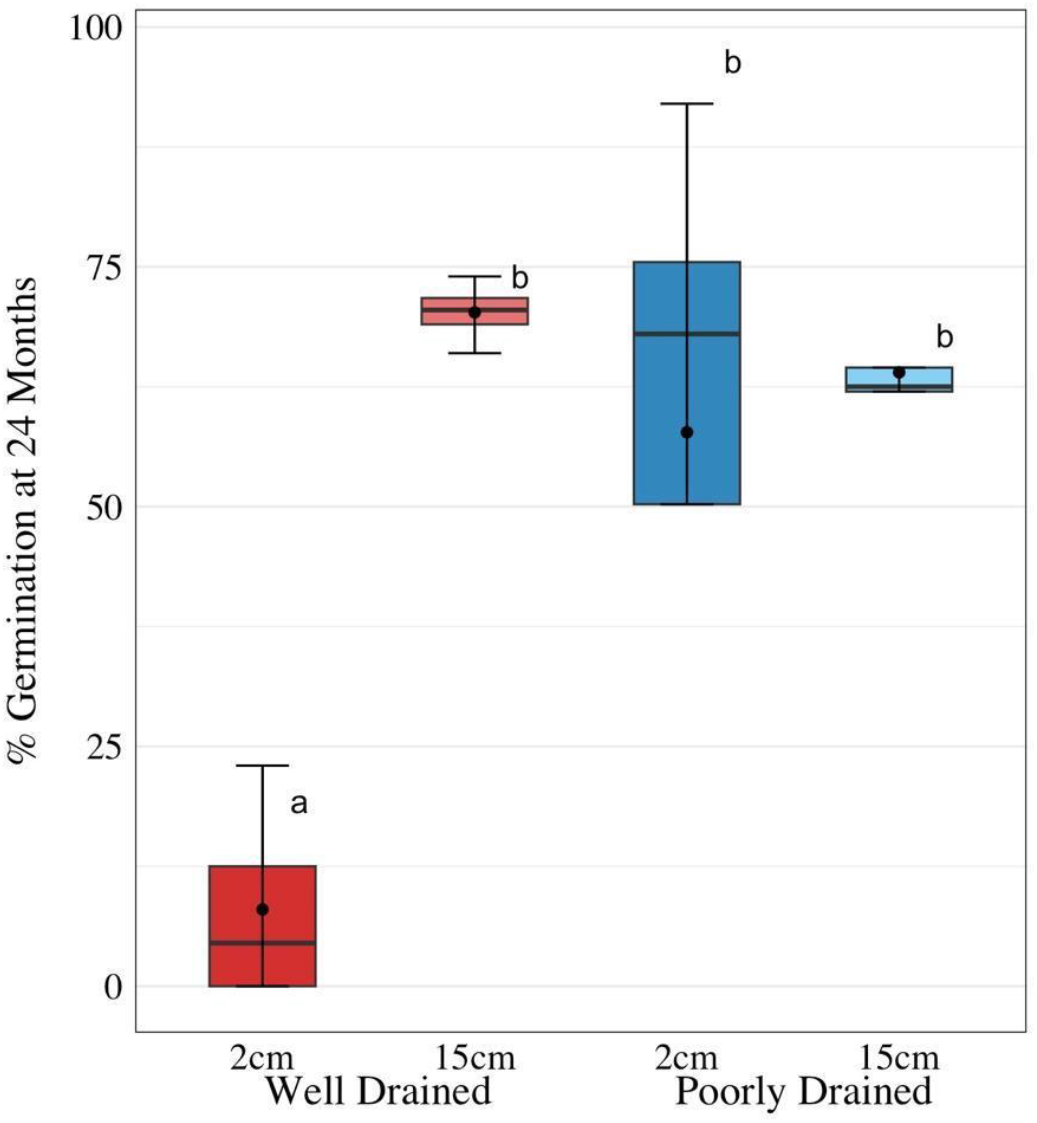
Percentage of seed germination averaged across all reps for black seeded ‘ARV1’. Seed burial depth significantly impacted germination percentages in well drained fields while, poorly-drained field conditions were not significantly impacted by burial depth.

Higher than average rainfall in August and September of 2022 and July and August of 2023 may have contributed to maintaining soil moisture throughout the typically drier months in Illinois in the poorly drained soil at the 15 cm depth (Table 2). The burial depth of wild pennycress at the 15 cm depth in the poorly-drained location showed a significant increase in seed viability compared to the other depths and locations (Figure 4). The increase in viability of the seed at the 15 cm depth of the poorly-drained field is likely due to diminished fluctuations in soil temperature and moisture. In addition, the wild thick seed coat of ‘ARV1’ would provide an extra layer of protection that would lead to seed that is more resilient to these fluctuations and remains viable longer.

**Table 2.**
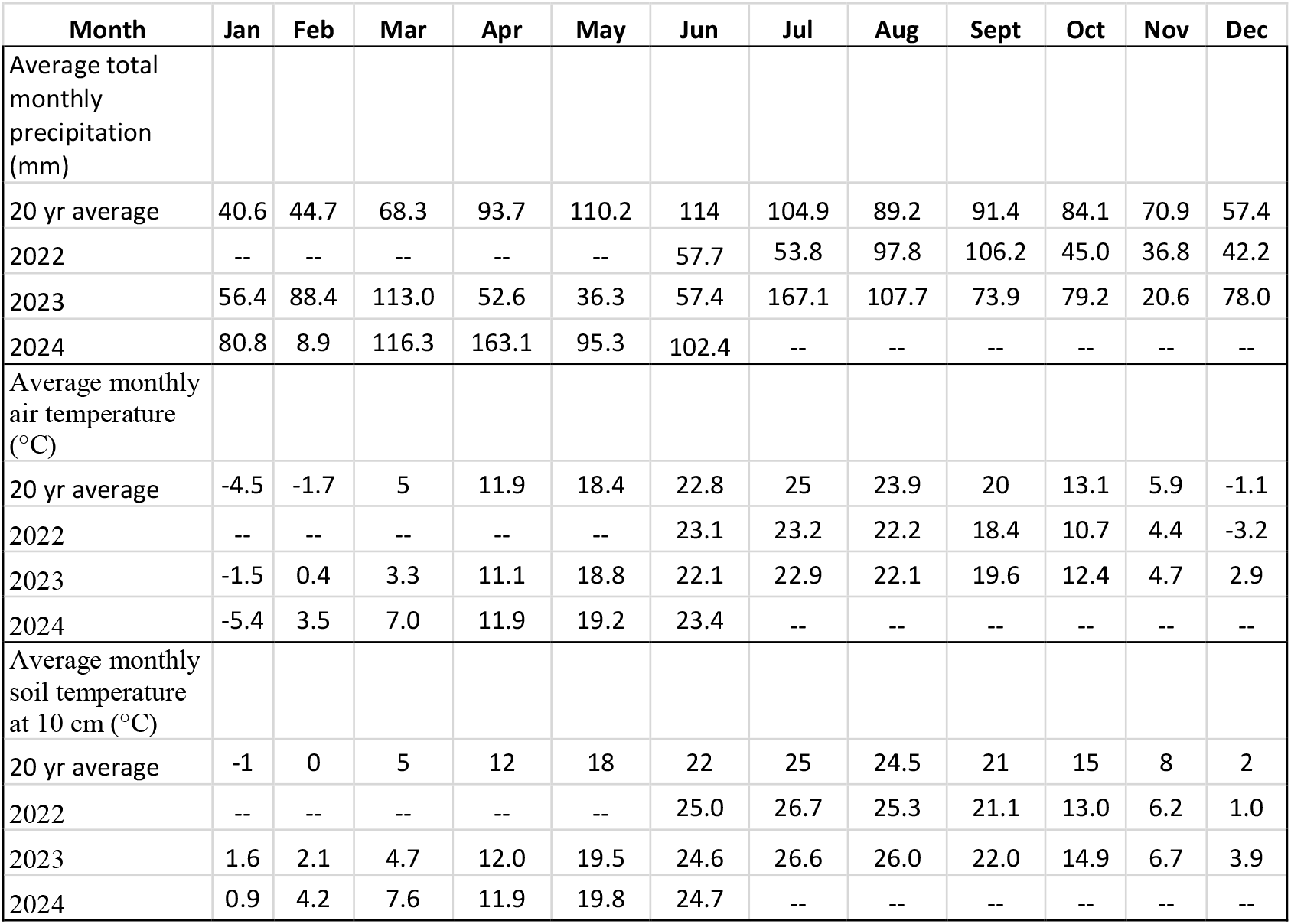
Comparison of monthly weather data for air temperature, soil temperature, and moisture at 10 cm to 20-year averages.

| Month | Jan | Feb | Mar | Apr | May | Jun | Jul | Aug | Sept | Oct | Nov | Dec |
| --- | --- | --- | --- | --- | --- | --- | --- | --- | --- | --- | --- | --- |
| Average total monthly precipitation (mm) |  |  |  |  |  |  |  |  |  |  |  |  |
| 20 yr average | 40.6 | 44.7 | 68.3 | 93.7 | 110.2 | 114 | 104.9 | 89.2 | 91.4 | 84.1 | 70.9 | 57.4 |
| 2022 | -- | -- | -- | -- | -- | 57.7 | 53.8 | 97.8 | 106.2 | 45.0 | 36.8 | 42.2 |
| 2023 | 56.4 | 88.4 | 113.0 | 52.6 | 36.3 | 57.4 | 167.1 | 107.7 | 73.9 | 79.2 | 20.6 | 78.0 |
| 2024 | 80.8 | 8.9 | 116.3 | 163.1 | 95.3 | 102.4 | -- | -- | -- | -- | -- | -- |
| Average monthly air temperature (°C) |  |  |  |  |  |  |  |  |  |  |  |  |
| 20 yr average | -4.5 | -1.7 | 5 | 11.9 | 18.4 | 22.8 | 25 | 23.9 | 20 | 13.1 | 5.9 | -1.1 |
| 2022 | -- | -- | -- | -- | -- | 23.1 | 23.2 | 22.2 | 18.4 | 10.7 | 4.4 | -3.2 |
| 2023 | -1.5 | 0.4 | 3.3 | 11.1 | 18.8 | 22.1 | 22.9 | 22.1 | 19.6 | 12.4 | 4.7 | 2.9 |
| 2024 | -5.4 | 3.5 | 7.0 | 11.9 | 19.2 | 23.4 | -- | -- | -- | -- | -- | -- |
| Average monthly soil temperature at 10 cm (°C) |  |  |  |  |  |  |  |  |  |  |  |  |
| 20 yr average | -1 | 0 | 5 | 12 | 18 | 22 | 25 | 24.5 | 21 | 15 | 8 | 2 |
| 2022 | -- | -- | -- | -- | -- | 25.0 | 26.7 | 25.3 | 21.1 | 13.0 | 6.2 | 1.0 |
| 2023 | 1.6 | 2.1 | 4.7 | 12.0 | 19.5 | 24.6 | 26.6 | 26.0 | 22.0 | 14.9 | 6.7 | 3.9 |
| 2024 | 0.9 | 4.2 | 7.6 | 11.9 | 19.8 | 24.7 | -- | -- | -- | -- | -- | -- |

## Conclusions

Golden gene-edited pennycress germination rates and seed viability diminished substantially in the first 2 months after burial with no viable seed remaining after 12 months of burial. In comparison, the wild type ‘ARV1’ had diminished germination rates over the 2 years but still had an average germination rate of over 50%. The golden lines were found to have a very low seed persistence and will not add to the weed seed bank when used in a crop rotation. The deletion of the *TT8* gene has greatly improved the domestication of pennycress and removed all concerns of seed carryover in production fields.

